# Heat-shock inducible clonal analysis reveals the stepwise establishment of cell fates in the rice stem

**DOI:** 10.1101/2023.04.26.538460

**Authors:** Katsutoshi Tsuda, Akiteru Maeno, Ken-Ichi Nonomura

## Abstract

The stem, consisting of nodes and internodes, is one of the major organs in seed plants. In contrast to other organs, however, processes of stem development remain elusive, especially when nodes and internodes are initiated. By introducing an intron into the *Cre* recombinase gene, we established a heat-shock inducible clonal analysis system in a single binary vector and applied it to the stem in the flag leaf phytomer of rice. With detailed characterizations of stem development, we show that cell fate acquisition for each domain of the stem occurs stepwise. Cell fates for a single phytomer and the foot (non-elongating domain at the stem base) were established in the shoot apical meristem by one plastochron before the leaf initiation. The fate acquisition for the node occurred just before the leaf initiation, separating cell lineages for leaves and stems. Subsequently, fates for the axillary bud were established in early leaf primordia. Finally, cells committed to the internode emerged from, at most, a few tiers of cells when the stem epidermis was at the 12∼25 celled stage. Thus, the internode is the last part of the stem whose cell fate is established. This study provides a groundwork to unveil underlying molecular mechanisms in stem development and a useful tool for clonal analysis, which can be applied to various species.

## Introduction

The clonal analysis gives insights into cell lineages and cell fate acquisition during the development of multicellular organisms. In principle, cells are genetically labeled at a certain point of development with a cell-autonomous marker(s) for visualization, and the extent to which the resulting clonal sector continues among tissues or organs is investigated. Such information can be used to deduce the location and number of cells which give rise to the tissue/organ of interest or to examine whether a certain cell fate has been established or not at the point of clone induction. For example, in maize (*Zea mays*), plants heterozygous for genes involved in anthocyanin or chlorophyll pigmentation were X-ray irradiated to delete dominant or functional alleles at various time points during development, leading to the expression of recessive sectors with reduced pigmentation (Johri and Coe, 1983; Poethig et al., 1986; McDaniel and Poethig, 1988; Poethig and Szymkowiak, 1995; Johri and Coe, 1996). Alternatively, in plants homozygous for recessive alleles with DNA transposon insertions, spontaneous reversions to the wild type in the presence of autonomous factors were utilized (Dawe and Freeling, 1990). In this case, the revertant sectors with pigmentation can be visualized. By using these methods, numbers and locations of cells in the embryonic shoot meristem contributing to adult organs such as leaves, internodes, ears (axillary branches bearing female inflorescence in maize), and tassels (male inflorescence) have been estimated (Johri and Coe, 1983; Poethig et al., 1986; McDaniel and Poethig, 1988; Poethig and Szymkowiak, 1995; Johri and Coe, 1996). Another example also in maize revealed that male gametes in anthers originated from the subepidermal L2 layer of shoot meristems (Dawe and Freeling, 1990). Recently, a heat-inducible clonal analysis in *Arabidopsis thaliana* roots determined the origin of vascular cambium stem cells and revealed the function of neighboring xylem cells as an organizer for the cambium (Smetana et al., 2019). Thus, clonal analysis is a powerful method to address important questions in plant development.

The stem is one of the major organs in vascular plants, which supports aboveground organs and connects the entire body through vascular networks. The stem in seed plants consists of reiterations of nodes and internodes. The node is an attachment point of a leaf to the stem in which the longitudinal growth is quite limited, whereas the internode is a domain of the stem that greatly elongates to lift leaves for light capture (**Figure 1, A and B**). The extent of stem elongation determines plant height. Therefore, the stem has been an important target in crop breeding, as exemplified by dwarf mutations utilized in the Green Revolution in the 1960s (Ferrero-Serrano et al., 2019). Despite its importance, however, the developmental process of the stem remains poorly studied. This is contrasting to other major organs such as roots, leaves, and flowers in which detailed regulatory mechanisms have been extensively studied, and is possibly due to a lack of clear landmarks specific to each part of the stem, at least externally, in many species (Serrano-Mislata and Sablowski, 2018).

**Figure 1.**
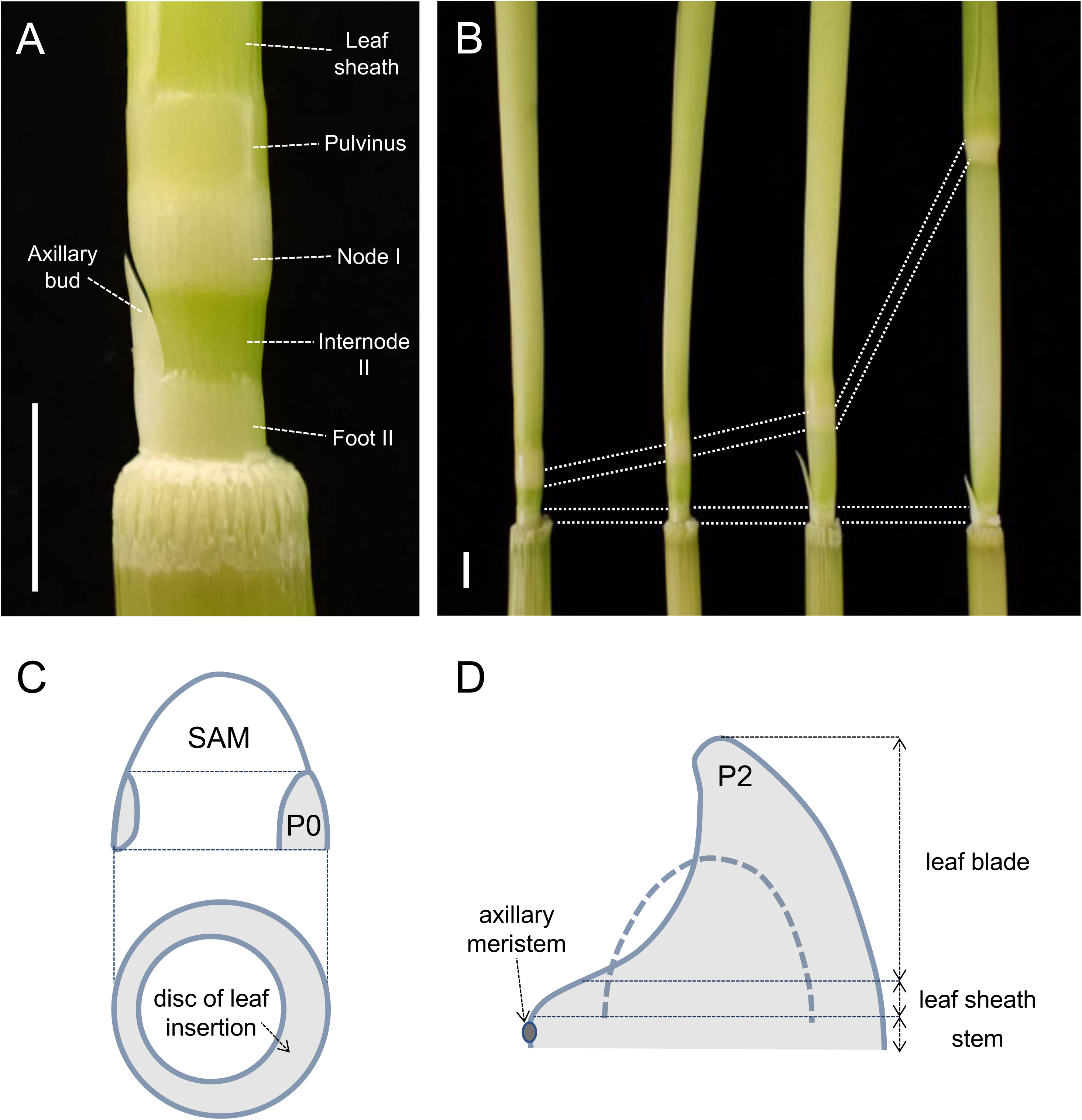
Illustration of the rice stem structure and a phytomer unit. (**A**) An immature stem of the flag leaf phytomer. **(B)** Stem samples at various stages of elongation. Note that only internodes elongate significantly. **(C)** A disc of leaf insertion (grey) from which a phytomer unit will form. This region corresponds to the P0 region where the initial down-regulation of *KNOX* genes occurs. **(D)** The P2 leaf primordium illustrating a phytomer unit which consists of a leaf blade at the top, a leaf sheath, an axillary bud, and a stem at the bottom. Scale bars are 5 mm.

The earliest event in stem development is the recruitment of leaf founder cells from the shoot apical meristem (SAM) (**Figure 1C**). Class I *knotted1-like homeobox* genes, which maintain an undifferentiated state of the SAM, are down-regulated in the P0 region (Jackson et al., 1994). This region corresponds to the disc of leaf insertion, which gives rise to a phytomer unit consisting of a leaf at the top, a node, an internode, and an axillary bud at the bottom (**Figure 1, C and D**) (Sharman, 1942). Based on histological observations, the upper and lower halves of the disc of insertion were suggested to develop into the leaf and the internode, respectively (Sharman, 1942).

Studies of clonal sectors induced in maize dry embryos or young seedlings showed that many sectors found in internodes started at the ear and extended into the leaf above (Johri and Coe, 1983; McDaniel and Poethig, 1988). Therefore, these organs often share a common cell lineage in early development, supporting the notion of a phytomer unit in grasses (Johri and Coe, 1983; McDaniel and Poethig, 1988). It was shown that sectors induced at one plastochron before leaf initiation were confined to a single phytomer but not to individual organs (i.e. leaf blades, sheathes, nodes, internodes, and axillary buds) (Poethig and Szymkowiak, 1995). Another study also in maize suggested that internodes remain a single or a few tiers of L1 cells after several plastochrons from their initiation (Johri and Coe, 1996). Thus, the cell fate for a single phytomer has already been established in the SAM just prior to the leaf initiation, but those for individual organs are likely to be specified later. It is still unknown, however, when and in what order the cell fates for individual organs, especially for the node and the internode, are established.

To address this question, a detailed analysis of clonal sectors induced at various time points of a certain phytomer development is important. In rice, the node and internode pattern is conspicuous due to extensive internode elongation after the reproductive transition (**Figure 1, A and B**). This transition is tightly controlled by the critical day length (Itoh et al., 2010), therefore, the flag leaf and accompanying stem with extensive elongation can be artificially induced by reducing the day length. Thus, rice can be a good model for studying stem development, and we aimed to establish a clonal analysis system in rice. The clonal analysis for vascular cambium in *Arabidopsis* mentioned above utilized a heat-inducible promoter and the Cre recombinase from the P1 bacteriophage to activate a GUS reporter (Smetana et al., 2019). In this study, we adopted this system with several modifications for rice and established a faithful induction of clonal sectors upon heat shock treatments. Using this system, we investigated the temporal order of cell fate establishment in the flag leaf phytomer in rice.

## Result

### Development of a clonal analysis system for rice in a single binary vector

The system of heat-shock inducible clonal analysis, originally developed in *Arabidopsis,* consisted of two binary vectors; one containing a heat shock-inducible promoter and the gene encoding Cre recombinase fused to CYCB1;1 destruction box (dBox) (hereafter called pHS_Cre), and another possessing the *35S* promoter, a roadblock (*loxP-tpCRT1-loxP*), and the *beta-glucuronidase* (*GUS*) gene (p35S_lox_GUS) (Smetana et al., 2019). The presence of dBox in the former vector results in the proteolysis of Cre proteins at the M phase in the cell cycle, avoiding a carryover of the protein. Upon heat shock treatments, the Cre recombinase removes the roadblock by recombining two *loxP* sites and allows the expression of *GUS* reporter. Because these recombination events occur by chance, the *GUS* reporter is activated in certain cells randomly, and such cells will generate clonal GUS-positive sectors.

To save time and effort due to two rounds of transformation, we aimed to combine the components into a single binary vector (**Figure 2, A and B**). It will be significant, especially in crop species with longer life cycles. Besides, we replaced the *35S* promoter with the maize *ubiquitin* (*UBQ*) promoter because we previously found that the *35S* promoter is often silenced and the *UBQ* promoter shows much more stable activity in rice (Tsuda et al., 2022). We also inserted a GFP coding sequence at the C terminus of GUS to allow non-destructive monitoring of spontaneous reporter activation. Initially, we tried to transfer this *proUBQ-loxP-tpCRT1-loxP-GUS-GFP* fragment into the *Cre*-containing binary vector, but it was unsuccessful. When we digested the resultant plasmid, the band pattern indicated that the plasmid lacked the *tpCRT1* roadblock (**Figure 2C**). This is possibly due to the misexpression of Cre recombinase in *Escherichia coli* and unwanted *loxP* recombination. Therefore, we inserted the first intron of the castor bean catalase gene *cat-1* into the Cre coding region. This intron is widely used in the intron-*GUS* reporter gene in binary vectors (Tanaka et al., 1990). As we expected, the resultant plasmid pHS_iCre_LGG_ver.1 (LGG stands for lox, <u>G</u>US and <u>G</u>FP) showed a predicted band pattern after restriction digestion, indicating that the introduction of the intron stabilized the plasmid structure (**Figure 2C**).

**Figure 2.**
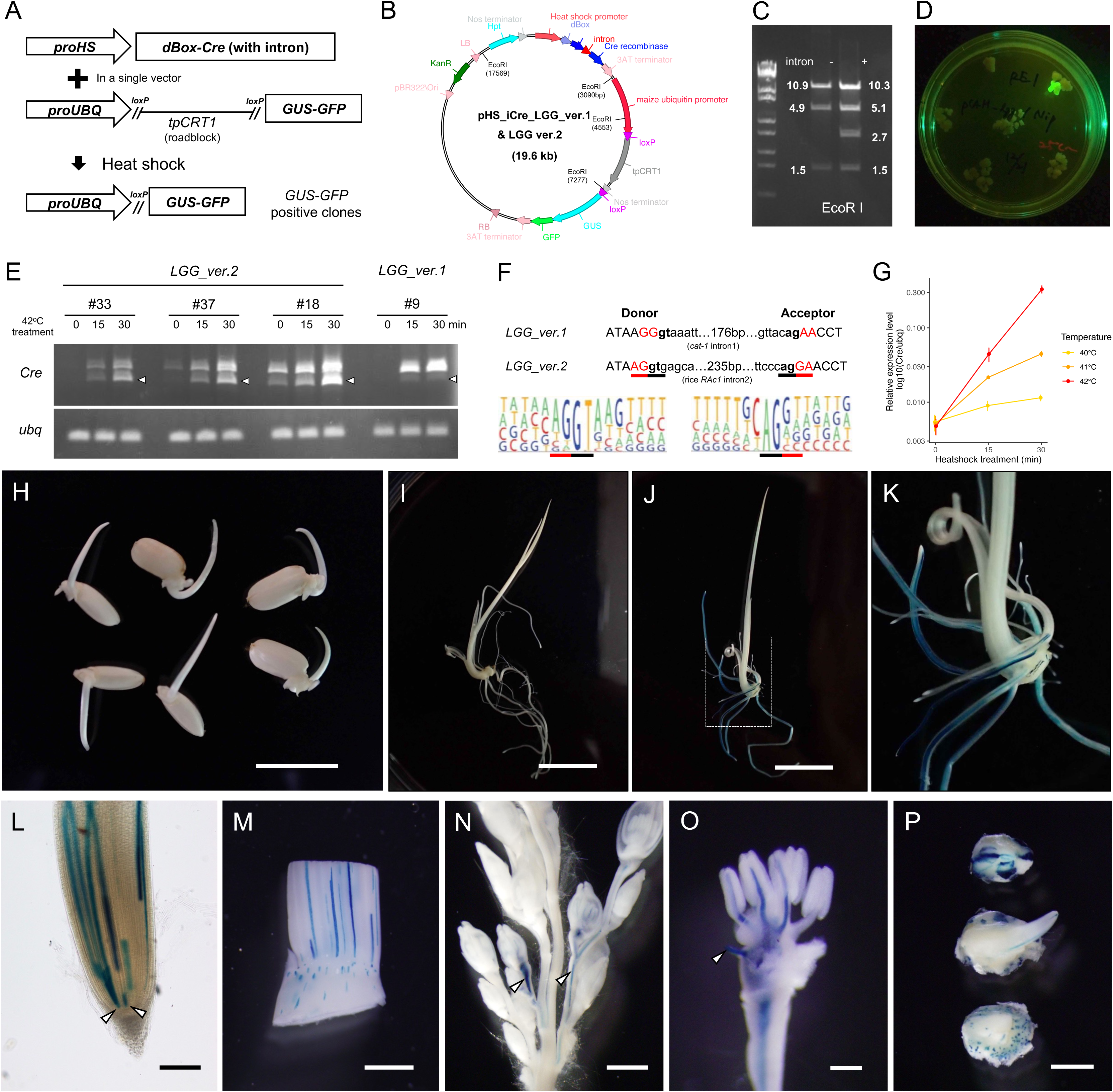
Development of a clonal analysis system in rice. (**A**) A schematic representation of this system. **(B)** A map of the components equipped in the vectors. EcoRI sites are indicated inside the circle. **(C)** EcoRI restriction digestion fragments of the plasmid with or without the *cat-1* intron. Note that the fragment at 2.7 kb corresponding to the *tpCRT1* roadblock is missing in the absence of the intron. **(D)** A spontaneous GUS-GFP reporter activation monitored by GFP. This example is from a single case observed in *LGG_ver.1*. **(E)** RT-PCR for intron-*Cre* genes. Arrowheads indicate the spliced form. Rice *ubq* was used as an internal control. **(F)** Comparison of the flanking sequences of the intron. DNA sequences of the first and second versions were indicated above the pictograms showing consensus sequences of the splicing donor and acceptor sites in the rice genome. Black and red underbars represent less variable dinucleotides in the intron and flanking exons, respectively. Pictograms are from Campbell *et al*. 2006. **(G)** qRT-PCR for the *Cre* during heat shock treatments. Error bars are standard deviations of three biological replicates. **(H)** Germinating seedlings at 3 DAG without induction. Note that there is no GUS staining. Two individuals of *LGG_ver.2* #33, 37, and #18 (from left to right) were stained for GUS. **(I)** A germinating seedling of *LGG_ver.2* #33 at 6 DAG without induction. There is no GUS staining. (J and K) A germinating seedling of *LGG_ver.2* #33 at 6 DAG with induction at 42°C for 30min. A dashed box in (J) indicates the region magnified in (K). **(L)** A root tip with multiple GUS sectors. Arrowheads indicate putative GUS sectors induced in stem cells. **(M)** GUS sectors induced in the shoot apex. Longitudinal cell files are conspicuous in the leaf sheath, whereas sectors induced in the non-elongating vegetative stem are confined to small regions. **(N)** Two inflorescence sectors (arrowheads) induced at the reproductive transition (+5 SD). **(O)** A sector in the spikelet, showing that three anthers share a common vascular cell lineage. An arrowhead indicates a vascular bundle entered into the lemma (removed). **(P)** Embryonic sectors induced during development and stained at 2 DAG. Scale bars are 5 mm in (H), 1 cm in (I) and (J), 200 µm in (L) and (O), 1 mm in (M), (N) and (P).

**Figure 3.**
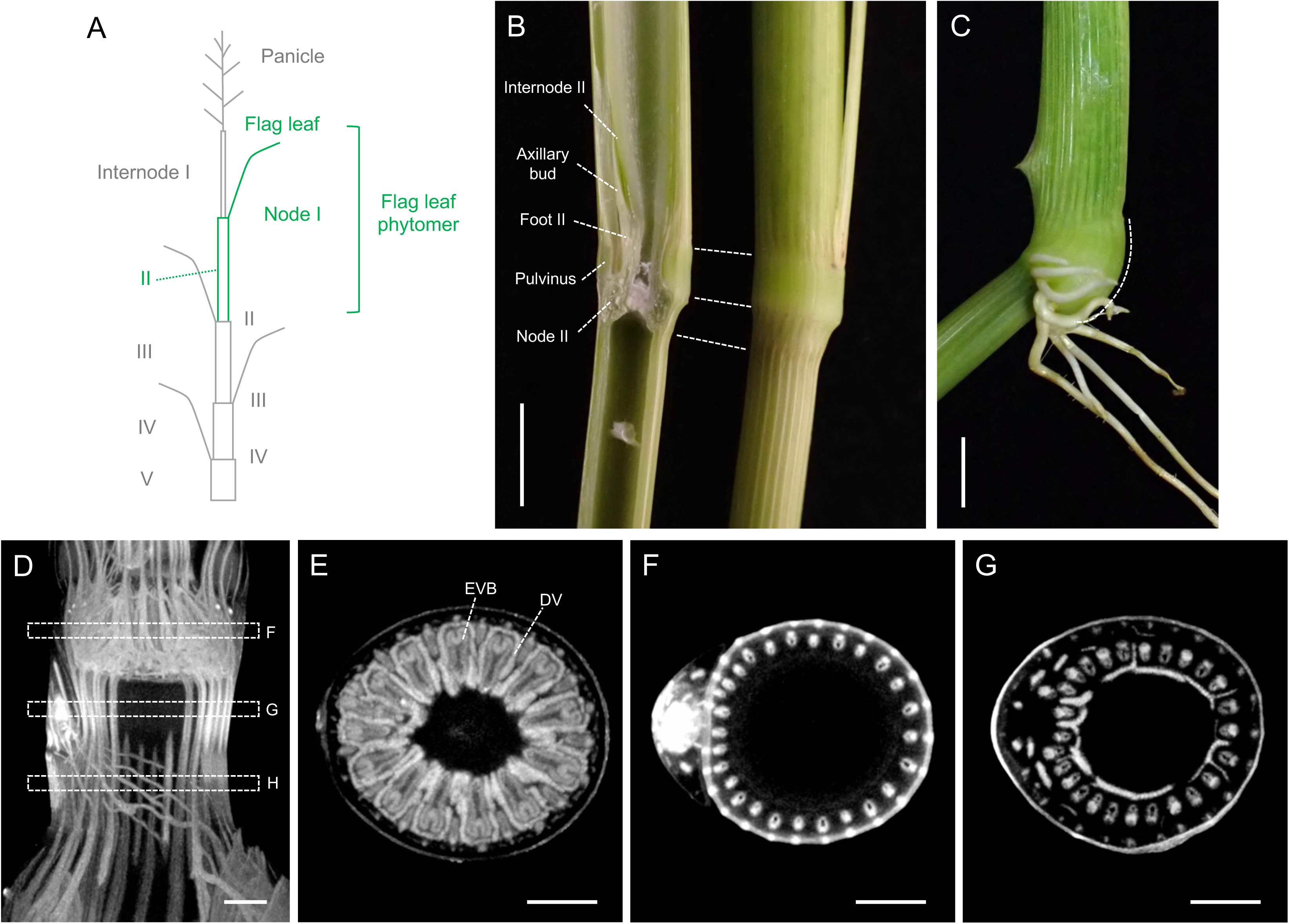
The structure of the rice stem. (**A**) The structure of rice plants and the numbering of each stem. Numbers for nodes and internodes are shown on the right and left, respectively. The flag leaf phytomer focused on in this study is colored green. **(B)** A vertical cut of the stem. Note that a central lacuna is continuous from internode II to foot II. **(C)** Node-specific structures. The pulvinus, whose bottom part expands (indicated by a dashed line), bends the stem in response to gravity. Crown roots had been initiated from the node. (D-G) Micro CT images representing internal structures of an immature stem of the flag leaf phytomer. (D) is a vertical section, and (E), (F) and (G) are horizontal sections at node I, internode II and foot II, respectively. Regions shown in (E) to (G) are indicated with dashed boxes in (D). EVB, enlarged vascular bundles; DV, diffuse vein. Scale bars are 5 mm in (B), (C), and 500 µm in (D)-(G).

### The initial version of the intron-*Cre* gene had a low splicing efficiency

We introduced pHS_iCre_LGG_ver.1 into rice calli (hereafter we call the transgenic plants *LGG_ver.1*) and monitored spontaneous reporter activation by observing GFP fluorescence (**Figure 2D**). Spontaneous activation in *LGG_ver.1* was rare (only one in 34 independent transgenic T0 calli). Next, we regenerated transgenic shoots from these calli and tested the induction rate of GUS sectors. Because it has been reported that a heat shock treatment at 42°C induces the expression of heat shock protein genes in rice (Hu et al., 2009; Zou et al., 2009), we treated *LGG_ver.1* regenerated plants at this temperature for 30 minutes. Among 15 independent transgenic T0 lines that we examined, only 9 sectors in 5 lines were found. Thus, even in the lines capable of induction, the induction rate was very low: one or two GUS sectors per plant. To determine a possible cause of this low induction rate, we checked the *Cre* gene expression by RT-PCR. We found that the spliced form of the *Cre* gene product was very faint, and the majority was in an unspliced form (**Figure 2E**). Thus, *LGG_ver.1* had a very low induction rate, possibly due to inefficient splicing of the intron introduced into the *Cre* gene.

### Adjusting the intron structure improved the splicing efficiency and sector induction rates

To improve the splicing efficiency, we compared the sequences around the splicing donor and acceptor sites of the inserted intron with those of the consensus in the rice genome (**Figure 2F**) (Campbell et al., 2006). Although the dinucleotides at the donor (GT) and acceptor (AG) sites were the same as the consensus, outside sequences in flanking exons differed (red underbars in **Figure 2F**). The upstream exons adjacent to the donor site in the consensus frequently end with a dinucleotide “AG”, whereas our case had “GG”. In addition, the downstream exons in the consensus frequently start with “G”, but our case did with “A”. This comparison suggested that the sequences around the intron insertion site were not optimal for efficient splicing in rice. It was also possible that the internal sequence of this *cat-1* intron was not suitable for efficient splicing in this specific case.

Based on these considerations, we shifted the intron insertion site 1 bp to the 5’ side to match the flanking exon sequences with those of consensus in the rice genome (**Figure 2F**). We also replaced the intron of *cat-1* gene with that of the rice actin gene, *RAc1*, which is highly and constitutively expressed (McElroy et al., 1990). We named this second version pHS_iCre_LGG_ver.2 and introduced it into rice calli. The frequency of the spontaneous reporter activation was similarly low as the first version (two in 46 independent transgenic T0 calli). Importantly, among the 30 T0 lines which were successfully regenerated, 27 showed the induction of multiple GUS-positive sectors after the heat shock treatment. Thus, the adjustment in the intron structure improved the splicing efficiency of the intron-*Cre* gene and enabled a successful induction of GUS sectors upon heat shock treatments.

### Testing the induction conditions for *Cre* gene expression and GUS sectors

To test the performance of this system, we selected three independent lines and examined the induction of the *Cre* gene in response to varying degrees of heat shock treatments. RT-PCR showed that the spliced *Cre* transcript in *LGG_ver.2* was induced upon heat shock, although we still detected a significant amount of the unspliced form (**Figure 2E**). In line #33, the *Cre* transcript was undetectable before induction, and the induction level was likely to be the lowest among the three. Upon heat shock treatments, the transcript level increased in the first 15 minutes, and the level further increased with longer incubation and/or at higher temperatures (**Figure 2, E and G**). Similar tendencies were found in the other two lines (#37 and #18), although they had leaky and higher expression levels (**Figure 2E**).

Next, we examined the efficiency of sector induction under various conditions (**Table 1**). We treated germinating seedlings 3 days after germination (DAG) and stained GUS sectors at 6 DAG. We counted the numbers of seedlings and roots (in both seminal and crown roots) with GUS sectors to evaluate the induction frequency. Importantly, without induction, germinating seedlings of these three lines at T2 generation showed no GUS staining, indicating that this system was kept uninduced through two rounds of generations under greenhouse conditions (**Figure 2, H and I, Table 1**). Incubation at 40 °C for 15 minutes induced no GUS sector. A small number of sectors was first observed after the treatment at 40 °C for 30 minutes (15.4 % of total plants and 3.7 % of total roots, **Table 1**). The induction rate significantly increased at 41°C; nearly half of the plants and a quarter of the roots generated GUS sectors after 15 minutes, and these rates were further increased after 30 minutes. At 42 °C, GUS sectors were observed in 75% of plants and 50% of roots after 15 minutes, and all individuals and 79.2 % of total roots had at least one sector after 30 minutes (**Figure 2, J and K, Table 1**). In line #18, which showed the highest *Cre* expression level, the induction rate was even higher (**Table 1**). Thus, the *Cre* expression levels and GUS-sector induction rates correlated. In contrast, the induction rate was very low in *LGG_ver.1*, indicating that the improvement of the intron was essential for efficient induction (**Table 1**).

**Table 1.** GUS sector induction rate in germinating seedlings.

| Line | Heat shock treatments | Number of plants with sectors | Total plants | Plants with sectors / total (%) | Number of roots with sectors | Total roots | Roots with sectors / total (%) |
| --- | --- | --- | --- | --- | --- | --- | --- |
| *LGG_ver.2* #33 | untreated | 0 | 14 | 0.0 | 0 | 73 | 0.0 |
|  | 40°C 15min | 0 | 12 | 0.0 | 0 | 58 | 0.0 |
|  | 40°C 30min | 2 | 13 | 15.4 | 2 | 54 | 3.7 |
|  | 41°C 15min | 7 | 13 | 53.8 | 14 | 61 | 23.0 |
|  | 41°C 30min | 10 | 13 | 76.9 | 21 | 59 | 35.6 |
|  | 42°C 15min | 9 | 12 | 75.0 | 29 | 60 | 48.3 |
|  | 42°C 30min | 14 | 14 | 100.0 | 61 | 77 | 79.2 |
| *LGG_ver.2* #37 | untreated | 0 | 7 | 0.0 | 0 | 42 | 0.0 |
|  | 42°C 15min | 4 | 7 | 57.1 | 8 | 34 | 23.5 |
|  | 42°C 30min | 7 | 7 | 100.0 | 41 | 48 | 85.4 |
| *LGG_ver.2* #18 | untreated | 0 | 5 | 0.0 | 0 | 29 | 0.0 |
|  | 42°C 15min | 5 | 5 | 100.0 | 23 | 25 | 92.0 |
|  | 42°C 30min | 5 | 5 | 100.0 | 27 | 28 | 96.4 |
| *LGG_ver.1* #9 | untreated | 0 | 9 | 0.0 | 0 | 53 | 0.0 |
|  | 42°C 15min | 1 | 8 | 12.5 | 1 | 45 | 2.2 |
|  | 42°C 30min | 2 | 8 | 25.0 | 5 | 50 | 10.0 |

### Sector inducibility in various tissues and organs

Next, we examined whether GUS sectors could be induced in various tissues and organs using *LGG_ver.2* #33 at T2 generation. In crown roots, as shown earlier, longitudinal cell files of GUS sectors were often observed (**Figure 2, J and K**). By closely examining their root apices, we identified sectors induced in the putative stem cell regions (**Figure 2L**). In the vegetative shoots, longitudinal cell files and small patches of GUS sectors were found in the young leaf sheath and stem, respectively (**Figure 2M, Supplemental Table 1**). Clonal analyses described in the following sections also showed that GUS sectors could be induced in the SAM during the reproductive transition (**Figures 4 and 5**). Furthermore, sectors could also be induced in developing panicles and embryos (**Figure 2, N-P**). Thus, these observations proved the utility of this system in studying various tissues and organs in rice.

**Figure 4.**
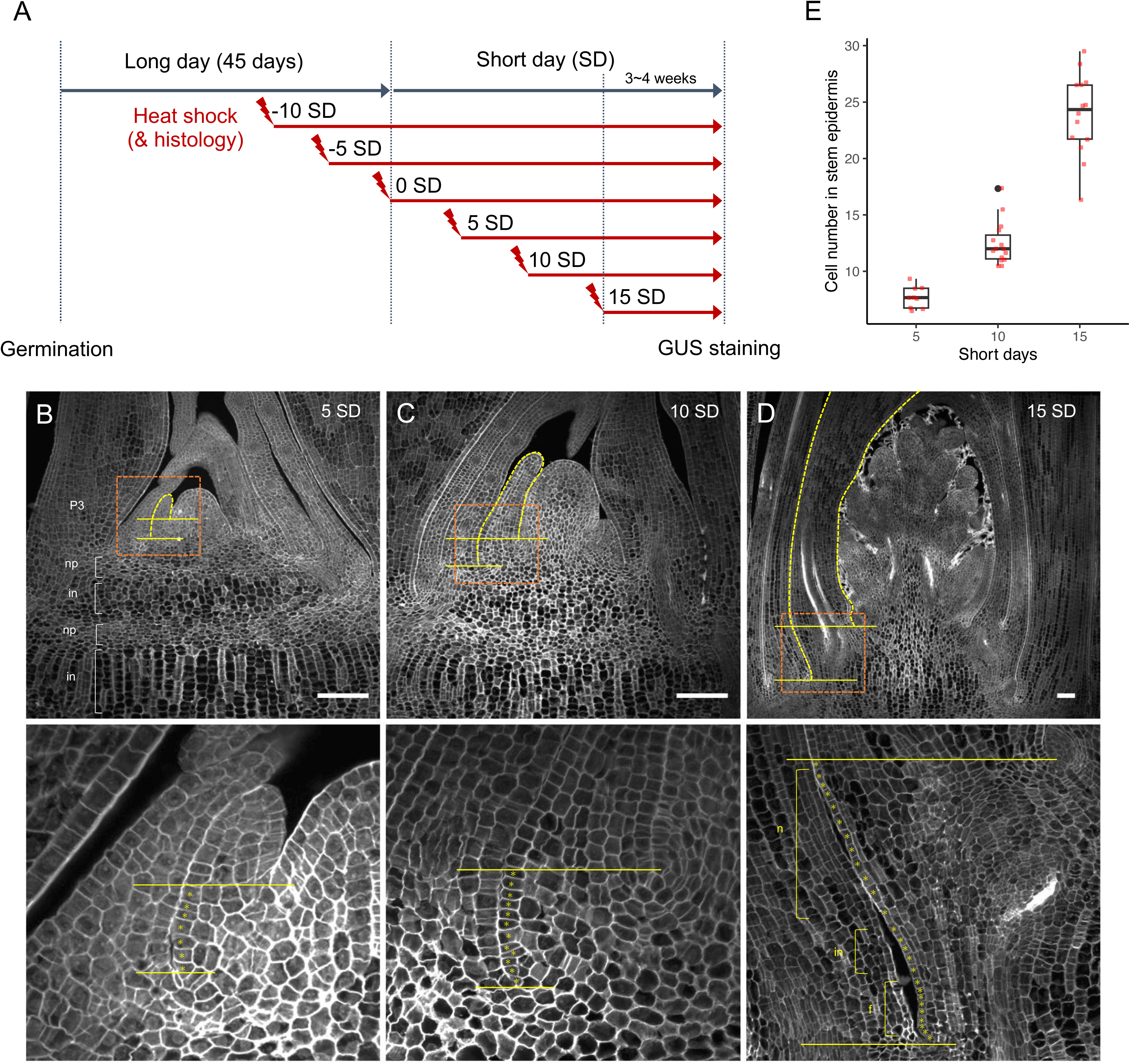
Histological observation of stem development. (**A**) A schematic representation of inducing the reproductive transition and heat shock treatments during stem development. (B-D) Confocal images of the shoot apex sections at +5 SD in (B), +10 SD in (C), and +15 SD in (D). Yellow horizontal lines indicate the bottom of leaf primordia to show young stem regions where epidermal cell numbers were counted. Yellow dashed lines indicate flag leaf primordia, and orange dashed boxes are approximate regions magnified in the panels below. Magnified images with clear cell boundaries are taken from sections different from those in the top panels. np, nodal plate; in, internode; n, node; f, foot. Asterisks in lower panels represent individual cells in the epidermis. Scale bars represent 100 µm. (E) Epidermal cell numbers of developing stem counted in tissue sections at +5, +10, and +15 SD. Red circles represent each sample.

**Figure 5.**
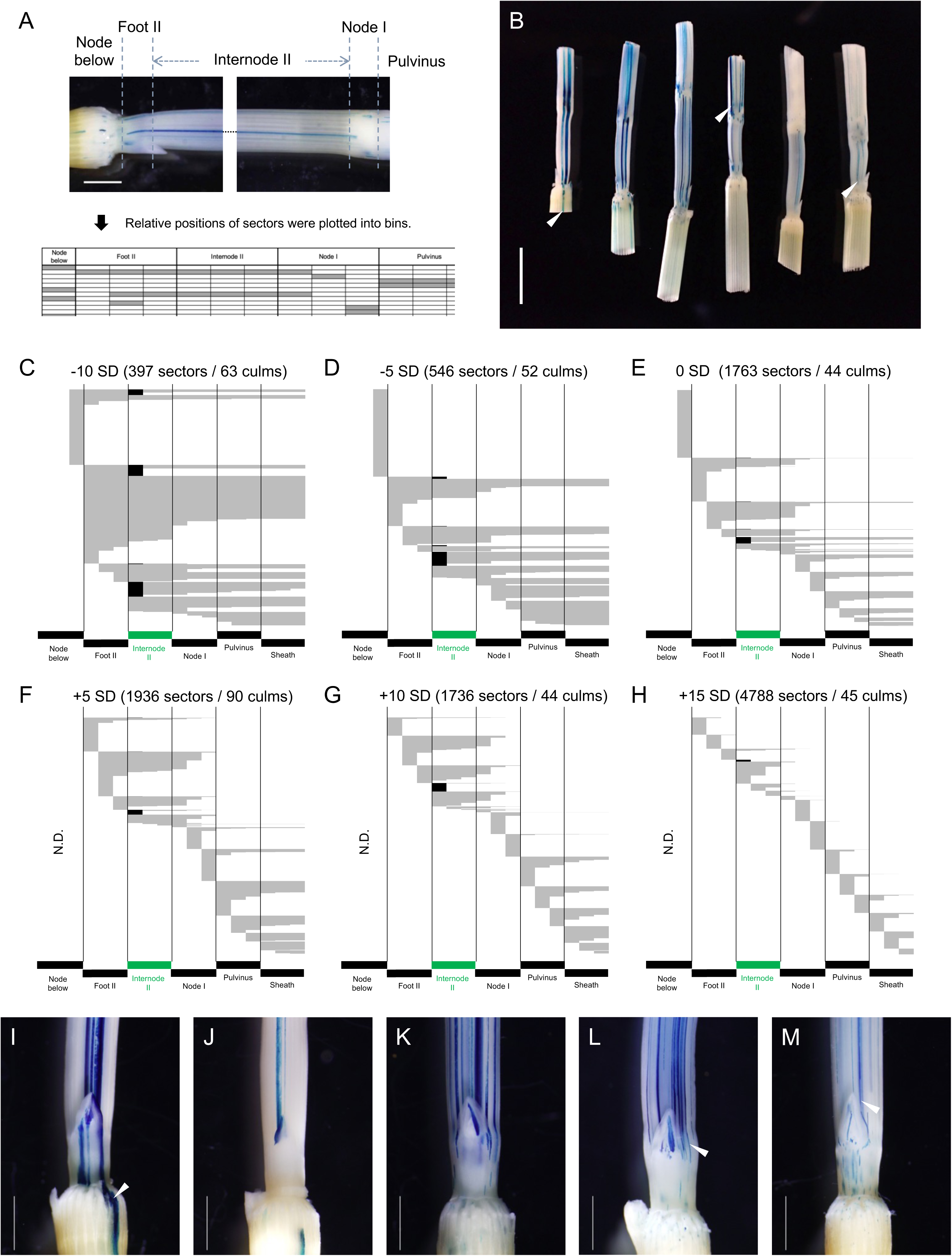
GUS sectors induced in the stem of the flag leaf phytomer. (**A**) GUS sectors and their assignment to each bin. Note that each domain of the phytomer is divided into three bins and the relative positions of sectors were assigned to these bins. (B) Representative GUS sectors induced at –10, –5, 0, +5, +10, and +15 SD from left to right. (C-H) The extent of GUS sectors induced at –10 to +15 SD in (C) to (H), respectively. Each sector is recorded in each row of the plot. Sectors found in axillary buds are colored black. The numbers of sectors and culms examined are shown at the top. (I-M) Examples of sectors that were induced in axillary buds at –10 to 0 SD (in I and J) and at +5, +10, and +15 SD in (K), (L), and (M), respectively. Arrowheads in (B) and (I) to (M) indicate sectors larger than their neighbors, exemplifying the variabilities of sector sizes. Bars represent 5mm in (B) and 2 mm (I) to (M).

### Characterization of the stem structure in rice

Before conducting clonal analyses, we characterized the structure and development of the stem in the flag leaf phytomer (**Figures 1 and 3**). Upon reproductive transition, the SAM produces the flag leaf and turns into the inflorescence meristem. The internode, whose elongation is limited during the vegetative phase, starts rapid elongation simultaneously in common rice cultivars. Elongating internodes are numbered from the top; the uppermost internode beneath the panicle is internode I, and the second one beneath the flag leaf is internode II (**Figure 3A**) (Takane and Hoshikawa, 1993; Yamaji and Ma, 2014). Nodes are also numbered in a similar way; the node at the flag leaf insertion (i.e., beneath the internode I) is called node I, and the next node below is node II. The foot, which is a non-elongating domain at the bottom (explained later), was numbered in a similar way in this study; that in between internode II and node II was named foot II (**Figure 1A, 3B**). Thus, the flag leaf phytomer consists of the flag leaf, node I, internode II, an axillary bud, and foot II.

Nodes are externally conspicuous in rice and other grasses, possibly because leaves encircle the entire circumference of the stem (**Figure 1A, 3B**) (Tsuda et al., 2017; Yamaji and Ma, 2017). Internally, node-specific enlarged vascular bundles (EVBs) derived from a leaf at the corresponding node connect to surrounding vascular bundles such as diffuse and transit vascular bundles originating from upper stems and to nodal anastomosis at the bottom of nodes (**Fugure 3, D and E**) (Yamaji and Ma, 2017). This complex vascular network is important for solute exchange because EVBs and surrounding veins/tissues are the site of action for transporters of various minerals (Yamaji and Ma, 2017). Nodes can also exert gravitropism owing to the pulvinus located at the base of the leaf sheath and are the site of initiating crown roots (**Figure 3C**). In internodes, vascular bundles are narrow and arranged vertically, presumably optimized for rapid elongation (**Figure 3, D and F**). Internode elongation is important to lift leaves and panicles for light capture and pollination, respectively. Thus, nodes and internodes have distinct but essential roles for survival. In rice, there is another non-elongating portion beneath internodes, which has not been well characterized yet (**Figure 1, A and B**). We recently named this region “foot” (Tanaka et al., 2023). Although the foot shares a central cavity with the above internode at maturity (**Figure 3B**), it is a chlorophyll-less and non-elongating tissue in which vascular bundles from the axillary bud connect to those of the stem (**Figure 1A, 3D, G**). The pulvinus surrounds the foot at the leaf base (**Figure 3B**). Therefore, it is appropriate to treat the foot as a structure distinct from the internode above.

### Histological observation of stem development

To characterize the process of stem development in rice, we conducted histological observations during the initiation of the flag leaf phytomer (**Figure 4** and **Supplemental Figure 1**). Because this reproductive transition is triggered by shifting day length from long to short day conditions in a photoperiod-sensitive cultivar Nipponbare (Itoh et al., 2010), the formation of this phytomer can be artificially induced (**Figure 4A**). After growing plants under long days for 45 days, the day length was shifted to short days. Because a new leaf is produced per 5 days on average in rice (Itoh et al., 1998), we took tissue samples every 5 days from 10 days before (–10 SD) until 15 days after the transition (+15 SD) from the top four shoots (**Figure 4A**).

At +5 SD, the flag leaf primordium had just been initiated (P1 primordium in **Figure 4B**). The number of epidermal cells in the future stem region of this primordium was around 7, and there was no sign of node and internode differentiation histologically (**Figure 4, B and E**). In the central region of the P3 stem, the putative node and internode organization was perceptible at this point (the upper “np” and “in” in **Figure 4B**), consistent with the previous observation (Kaufman P, 1959). Nodal plates consist of flattened and less ordered cells, whereas the putative internodes in the center form longitudinal files of elongating cells (**Figure 4B**). However, at the periphery of these stems, cells do not show such specialized organizations (**Supplemental Figure 1, A-C**). Hence, differentiation along the longitudinal axis remains unclear in the stem periphery. At +10 SD, the flag leaf becomes hood shaped and covers the enlarging inflorescence meristem (**Figure 4C** and **Supplemental Figure 1D**). The young stem beneath the flag leaf had ∼12 epidermal cells at this stage, and there was still no sign of differentiation (**Figure 4, C and E**). At +15 SD, when the inflorescence branching had started, the cell number in the stem epidermis reached ∼25 cells (**Figure 4, D and E**). At this stage, a putative organization of the node and internode pattern can be seen in the peripheral region (**Figure 4D, Supplemental Figure 1, F and G**). Subepidermal cells in putative node I and foot II are less ordered, whereas those in putative internode II are arranged in longitudinal files. This organization is more evident in the stem one phytomer below (**Supplemental Figure 1H**). Taking together, although the differentiations of putative nodes and internodes become distinct in early development in the central region of the stem, those at the periphery, which give rise to the main stem tissue at maturity, remain unclear until later stages.

### Induction of GUS sectors during the reproductive transition

To investigate the timing and the order of cell fate establishment in the stem, we applied our clonal analysis system to the stem development of the flag leaf phytomer. Here, we follow the notion of cell identity acquisition; it is not the lineage but the position of cells relative to their surroundings that determines cellular identities in many aspects of plant development (Poethig et al., 1986; McDaniel and Poethig, 1988; Steeves and Sussex, 1989; Scheres, 2001). If a certain sector was confined to a single domain (organ or tissue), we consider that cells within the sector did not proliferate beyond the domain boundaries and hence that the cell fates had been established at the point of induction.

Along with the growth scheme of the above-mentioned reproductive transition, we prepared six sample groups at intervals of 5 days (**Figure 4A**). The heat shock conditions were at 42℃ for 30 or 45 minutes because treatments of lower temperature or shorter time did not induce enough sectors in the stem (**Supplemental Table 2**). After the last treatment at +15 SD, we let *LGG_ver.2* #33 plants grow for additional 3∼4 weeks and harvested culms for GUS staining. Each sample group comprised the top 4 or 5 culms from at least 10 individuals. Because the internode II was much longer than other domains, we plotted the relative positions of GUS sectors into three bins for each domain (**Figure 5A**). Besides, as samples were cut at node II or internode III for the lower end and at the middle of the flag leaf sheath for the upper end, we did not deal with sectors found in these regions except when the continuity of sectors from the neighboring domains was considered.

### The earliest events were the establishment of cell fates for the flag leaf phytomer and foot II

The number of sectors ranged from 397 to 4788, depending on the timing of the heat shock treatment (**Figure 5, C-H**). Samples induced at later time points tended to have more sectors because the number of cells in the tissues of interest increased as the plant grew. Overall, GUS sectors initially spanned broadly but were gradually confined to narrower regions of the stem (**Figure 5, B-H**, and **Table 2, 3**).

**Table 2.** The number of sectors extended for multiple tissue types (TT).

| Stage | Two phytomers | Entire flag phytomer | Four TTs | Three TTs | Two TTs | Single TT | Total |
| --- | --- | --- | --- | --- | --- | --- | --- |
| -10 SD | 25 (8.7) | 94 (32.8) | 32 (11.1) | 32 (11.1) | 65 (22.6) | 39 (13.6) | 287 |
| -5 SD | 0 | 40 (11.8) | 35 (10.3) | 101(29.7) | 79 (23.2) | 85 (25.0) | 340 |
| 0 SD | 0 | 12 (1.0) | 31 (2.5) | 275 (22.0) | 240 (19.2) | 691 (55.3) | 1249 |
| +5 SD | 0 | 0 | 1 (0.1) | 267 (13.7) | 424 (21.7) | 1229 (64.5) | 1952 |
| +10 SD | 0 | 0 | 0 | 154 (8.6) | 357 (19.9) | 1284 (71.5) | 1795 |
| +15 SD | 0 | 0 | 0 | 14 (0.3) | 191 (4.0) | 4546 (95.7) | 4788 |

Numbers in parentheses are percentages by total sectors.
Sectors confined in node II were not considered.

In samples treated at –10 SD, most sectors (91.3%) were confined to the flag leaf phytomer (**Table 2** and **Figure 5C**). Among these, sectors spanning the entire flag leaf phytomer were the most prominent (32.8%), and those across multiple tissue types were also frequent (**Table 2**). This indicates that, at two plastochrons prior to the flag leaf initiation, the cell fates for this phytomer but not for specific organs had been largely established in the SAM. At significant frequencies, however, we observed sectors confined to the bottom of foot II (12.5%, **Table 3**), suggesting that foot II may be the earliest domain whose cell fate is established. Occasionally, we also observed sectors that spanned across two successive phytomers (8.7%, **Table 2**). This reflects the existence of cells whose fate had not been destined to a single phytomer in the sample population at a certain frequency.

**Table 3.** Sectors confined in a single tissue type (TT).

| Stage | Foot | Axillary bud | Internode | Node | Pulvinus | Sheath | Single TT<br>sectors | Total sectors |
| --- | --- | --- | --- | --- | --- | --- | --- | --- |
| -10 SD | 36 (12.5) | 3 (1.0) | 0 | 0 | 0 | 0 | 39 (13.6) | 287 |
| -5 SD | 68 (20.0) | 5 (1.5) | 0 | 10 (2.9) | 0 | 2 (0.6) | 85 (25.0) | 340 |
| 0 SD | 332 (26.6) | 11 (0.9) | 0 | 206 (16.5) | 112 (9.0) | 30 (2.4) | 691 (55.3) | 1249 |
| +5 SD | 487 (24.9) | 31 (1.6) | 0 | 394 (20.2) | 247 (12.7) | 101 (5.2) | 1260 (64.5) | 1952 |
| +10 SD | 270 (15.0) | 61 (3.4) | 50 (2.8) | 346 (19.3) | 349 (19.4) | 208 (11.6) | 1284 (71.5) | 1795 |
| +15 SD | 812 (17.0) | 37 (0.8) | 653 (13.6) | 1081 (22.6) | 1161 (24.2) | 839 (17.5) | 4583 (95.7) | 4788 |

Numbers in parentheses are percentages by total sectors.
Sectors confined in node II were not considered.

In samples treated at –5 SD, sectors that spanned across two phytomers disappeared, although 40.9% of the sectors extended both into the flag leaf sheath and the stem (**Figure 5D** and **Supplenental Table 3**). Therefore, the fate of the flag leaf phytomer had been completely established but has yet to be for specific organs. In addition, sectors confined to foot II became more evident (20%, **Table 3**), supporting the hypothesis that foot II is the earliest domain whose cell fate is established.

### Determination of nodal cell fates split the flag leaf and the subtending stem

In samples treated at 0 SD, sectors that extended for the entire phytomer almost disappeared (1%), and those confined either to the stem or the flag leaf became evident (88.8%, **Table 2, Supplemental Table 3** and **Figure 5E**). Therefore, cell lineages specifically contributing to the flag leaf or the stem had been largely established at the flank of the SAM, or in the P0 region. This divergence of cell lineages for the flag leaf and the stem was accompanied by the frequent emergence of sectors confined in node I or the pulvinus (25.5%, **Table 3**). These tendencies became more evident in the samples treated at +5 SD (**Figure 5F**). Therefore, node I and the associated leaf structure pulvinus may be second domains whose cell fate was established, splitting neighboring cell lineages into the flag leaf and subtending stem. It is noteworthy that, at 0 SD, sectors confined to the axillary bud and internode II were rare and absent, respectively; sectors found in axillary buds often extended to internode II, and those in the internode II always spanned either to the foot II or node I (**Figure 5, E and J**). These indicate that cell fates for the axillary bud and internode II had not been established at 0 SD.

### The internode is the last part of the stem whose cell fate was established

In contrast to earlier sample groups, sectors in axillary buds induced at +5 SD hardly extended into internode II anymore (**Figure 5, F, K-M**). The size of sectors in axillary buds gradually became smaller in samples treated later. These observations suggested that cells contributing to axillary buds had established their fates by the time the flag leaf primordium encircles the SAM (**Figure 4B**). In contrast, all sectors found in internode II spanned to neighboring tissues (**Figure 5, F and K**). Thus, the cell fate for internode II had yet to be established.

Sectors confined to internode II were first observed at a low frequency (2.8 %) in samples treated at +10 SD (**Figure 5G** and **Table 3**). In the sample group treated at +15 SD, most GUS sectors found in internode II (88.8%) did not extend into neighboring tissues anymore (**Figure 5H, Table 3** and **Supplemental Table 3**). Therefore, cell fate determination for the internode likely occurred between the 12-and 25-celled stages. Notably, the majority of these sectors in internode II spanned more than half of the length of internode II (**Figure 5H**), suggesting that the number of cells destined for internode II was still very few at +15 SD. These results showed that the internode II was the last tissue whose cell fate was determined in the flag leaf phytomer.

## Discussion

In this study, we established an efficient tool for clonal analysis in rice. Based on the detailed description of the stem development, we applied our tool to reveal the temporal order of cell fate determination in the stem tissues of the flag leaf phytomer. Because our system can be readily applied to any organs in rice examined so far, it will be a powerful tool to widen the knowledge of cell lineage and cell fate determination, which is still very limited in rice. Inserting an intron into the Cre coding region enabled us to unite the system components into a binary vector, allowing a rapid establishment of experimental lines through a single transformation. Moreover, because the heat shock treatment requires no chemical inducers such as dexamethasone or estradiol, it is harmless to researchers. It can be applied to a wide range of plant species with various sizes.

Spontaneous activation of GUS sectors was rare, and GUS sector induction strictly required heat shock treatments even with a leaky expression of the *Cre* gene. The unspliced form of the *Cre* transcript was still detected in the improved version *LGG_ver.2.* Therefore, it is possible that this incomplete splicing hindered the accumulation of the mature transcript above a certain threshold and hence might have served as a rate-limiting step for induction. In this scenario, the level of mature *Cre* transcripts in the leaky lines may be simply under this threshold before the heat shock. Spontaneous inductions of sectors that we observed in calli may be the cases in which the basal expression level of *Cre* exceeded this level. Although the exact mechanism for this requirement of heat shock is unclear, the insertion of an intron likely resulted in a faithful induction system.

The frequency of GUS-sector induction increased with higher temperatures and longer treatments. In germinating seedlings, efficient induction of GUS sectors was already observed at 41°C, and upregulation of *Cre* was observed within 15 minutes after the treatment. However, in the case of the flag leaf phytomer, the heat shock treatment at 42°C for 30 minutes or longer was required for an efficient induction (**Supplemental table 2**). This is probably due to the requirement of higher temperatures or a longer time for heat conduction to reach the shoot apices enclosed by adult leaves. Thus, the optimal condition for heat shock depends on how the tissue of interest is exposed. It is important to select the lowest temperature and the shortest duration to avoid undesired side effects such as growth retardation or tissue damage.

In every sample group, we observed significant variabilities in the extent that sectors extended. For example, most sectors induced at –10 SD were confined to the flag leaf phytomer, whereas some spanned to the neighboring node II. In axillary buds, although most sectors extended to neighboring organs until +5 SD, those confined to this organ were observed at low frequencies repeatedly (Table **4**). Even in a single sample, some sectors extended for more numbers of phytomers/organs/tissue types than their neighboring sectors (arrowheads in **Figure 5, B, I, L, M**). According to the studies in maize, there are two possible sources of these variabilities (Poethig et al., 1986; McDaniel and Poethig, 1988). First, there must be variances in developmental stages among samples at the point of induction. This is trivial but unavoidable. Second, more importantly, cells at similar positions in a certain tissue may contribute differently due to the variance in the frequency and orientation of cell division. Thus, even at similar positions, individual cells can vary in their final lineage sizes or identities, and the sectors confined to a single domain found in earlier samples may reflect the existence of cells that do not actively divide until late development.

Our experiments showed that the cell fates for distinct parts of the stem are determined stepwise depending on the tissue type (summarized in **Figure 6**). Around –10 SD when the leaf primordium two plastochrons before the flag leaf is initiating, the cell fate for the flag leaf phytomer is largely but incompletely established in the SAM, and it becomes complete 5 days later. This is consistent with the observation in maize, in which the fate for a single phytomer is established one plastochron before its initiation (Poethig and Szymkowiak, 1995). In addition, a fraction of the cell population had already been committed to the bottom part of foot II, indicating that the most bottom tiers of cells destined for the flag leaf phytomer in the SAM will be the first tissue whose cell fate is established. At the onset of the reproductive transition (0 SD), cell fates for the nodal tissue, including node I and the pulvinus, start being established at the flank of the SAM, leading to the divergence of cell lineages for the flag leaf or the stem. This result supports the hypothesis proposed by Sharman, in which the upper and lower halves of the disc of leaf insertion contribute to the leaf and the stem, respectively (Sharman, 1942). Subsequently, cells located beneath the overlapping margins of the leaf sheath become committed to the axillary bud early in the flag leaf primordium (+5 SD). It is not until +10 to +15 SD that the cell fate for internode II is established. The number of epidermal cells in the developing stem at this point ranges from 12 to 25, and the cell fate commitment for internode II likely occurs in a single to a few tiers of cells. This is consistent with the report in maize, in which internodes remain a single or a few tiers of L1 cells after several plastochrons from their initiation (Johri and Coe, 1996). Overall, cell fate commitment for non-elongating tissues occurs first, in which the foot is determined earlier than the node, and that for the internode takes place at the end.

**Figure 6.**
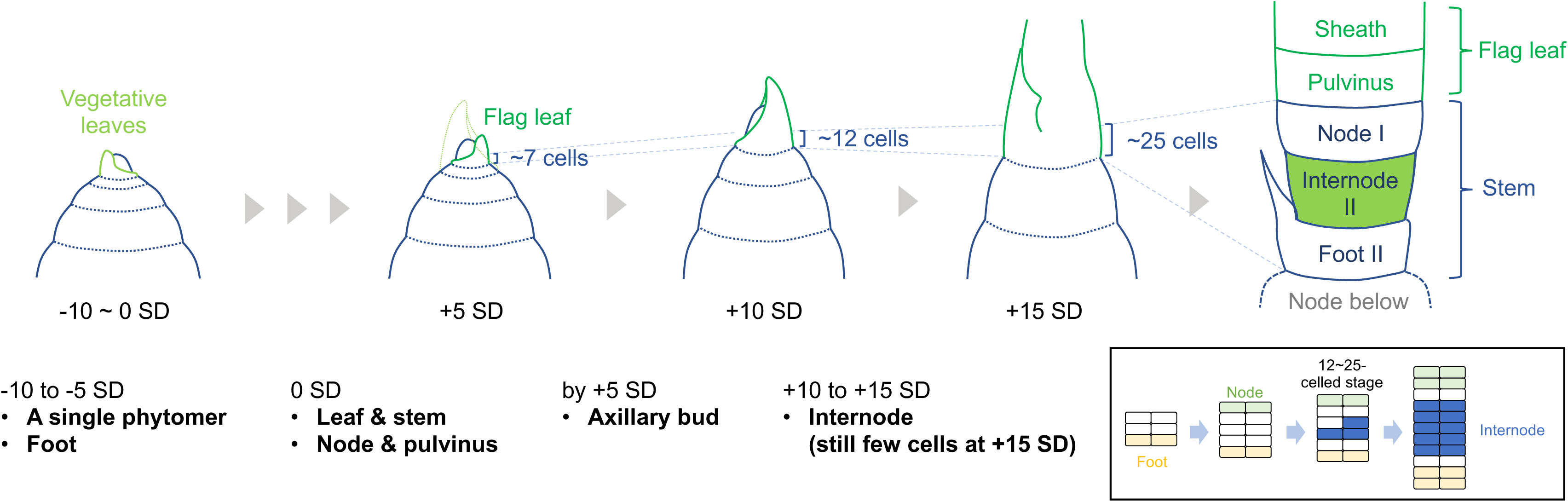
The temporal order of cell fate acquisition of the rice stem. The fate of the flag leaf phytomer is established in one to two plastochrons prior to the leaf initiation. Non-elongating tissues such as foot II, node I, and the pulvinus acquire their fate earlier than elongating tissues. Internode II is the last part of the stem whose cell fate is established. It is likely to originate from a very limited number of cells between node I and foot II (inset).

An intriguing question is how the cell/tissue identities along the vertical axis of the stem are determined. Our data suggested that the cell fates for the node and its associated pulvinus are established in the SAM before the initiation of flag leaf primordia, whereas that for the internode is determined in a very limited number of cells much later (inset in **Figure 6**). This is in clear contrast to the central region of the stem in which the putative node and internode patterns emerge almost simultaneously. The central regions in the internode and the foot form cavities or lacunae for gas exchange, possibly through programmed cell death (Steffens et al., 2011; Fujimoto et al., 2018), whereas this does not occur in the peripheral region. It is possible that distinct regulatory programs govern patterning events in the central and peripheral regions of the stem from the very early stage of development. Studies in deepwater rice have shown the presence of several activators and a repressor of internode elongation, which confer the regulated competency for internode growth depending on water levels, even in the vegetative phase (Hattori et al., 2009; Kuroha et al., 2018; Nagai et al., 2020). In the vegetative stem of common cultivars, internodes may be present but simply cannot elongate due to the repressor (Gomez-Ariza et al., 2019). Alternatively, internode initiation *per se* may be repressed in this phase. So far, molecular mechanisms involved in these regulations and those that specify the node and internode pattern are entirely unknown. The stepwise establishment of cell fates revealed here will be the foundation to unveil underlying mechanisms in stem development by providing clues regarding the timing of key events in future studies.

## Materials and Methods

### Plasmid construction

The original heat-shock inducible clonal analysis system reported by Smetana *et al*. consists of two binary vectors; one containing a heat shock-inducible promoter and the gene encoding Cre recombinase fused to CYCB1;1 destruction box (dBox) (called pHS_Cre in this study), and another possessing the *35S* promoter, a roadblock (*loxP-tpCRT1-loxP*) and a beta-glucuronidase (GUS) coding region (p35S_lox_GUS) (Smetana et al., 2019). To combine these components into a single binary vector, we mutated the original KpnI site in pHS_Cre and re-introduced a new KpnI site next to the SacI site by ligating annealed double-stranded oligonucleotides (KT1320-1321). This vector containing the neighboring SacI and KpnI cloning sites was named pHS_Cre2.

We also modified p35S_lox_GUS to achieve stable expression and non-destructive monitoring of spontaneous reporter activation in rice. To place the *loxP-tpCRT1-loxP-GUS* under the maize *ubiquitin* (*UBQ*) promoter, we subcloned a 4.6 kb HindIII fragment containing these elements from p35S_lox_GUS into pPUBn (Tsuda et al., 2022) and named this plasmid pUBQ_lox_GUS. Then we fused GFP to the C terminus of GUS through seamless cloning (NEBuilder, NEB) using primers KT1101, KT1102, KT1103, and KT1104. We named this plasmid pUBQ_LGG. The first intron sequence from the castor bean *cat-1* gene in the pBGH1 vector (Ito and Kurata, 2008) was introduced into pHS_Cre through NEBuilder using primer sets KT1387, KT1388, KT1389, and KT1390. The resultant plasmid was named pHS_iCre.

A 7.4kb SacI-KpnI fragment containing *pUBQ*-*loxP-tpCRT1-loxP-GUS-GFP* from pUBQ_LGG was subcloned into pHS_iCre, and this plasmid was named pHS_iCre_LGG_ver.1.

We replaced the *cat-1* intron with the second intron of the rice *Actin1* gene through NEBuilder using primers KT1491, KT1492, KT1493, and KT1494, and named the resultant plasmid pHS_iCre2. Finally, the 7.4kb SacI-KpnI fragment from pUBQ_LGG was subcloned into pHS_iCre2 and this plasmid was named pHS_iCre_LGG_ver.2.

DNA sequences amplified using PCR were confirmed by sequencing. Primers used in the plasmid construction are listed in **Supplemental Tabel 4**.

### Plant materials, transformation, and growth conditions

Transgenes were introduced into Nipponbare using *Agrobacterium*-mediated transformation as described previously (Toki et al., 2006). Hygromycin-resistant plants were transferred to soil in black vinyl pots (13.5 cm diameter) until maturity. Transgenic plants at subsequent generations were genotyped for the *hygromycin phosphotransferase* (*hpt*) gene using primers KT287 and KT288, and *hpt*-positive plants were grown in the growth chamber under the long-day (LD) condition (14-h light at 30°C and 10h dark at 25°C). At 45 days after germination (DAG), the day-night cycle was shifted to the short-day (SD) condition (10-h light at 30°C and 14-h dark at 25°C).

### Heat-shock treatments

To test heat-shock conditions, transgenic seedlings at 3 DAG were incubated in a water bath at 40, 41, and 42°C for 15 and 30 minutes. After the heat shock, plants were grown for 3 days in the growth chamber until GUS staining.

For sector induction in the internode II, plants at 35, 40, 45, 50, 55, and 60 DAG (for –10, –5, 0, +5, +10, and +15 SD treatments, respectively) were incubated in a water bath at 41 or 42°C for 30 or 45 minutes. To avoid the temporal drop of water temperature at the beginning of the heat shock, up to 3 pots with soil were incubated in 35 L of water at once, and the water temperature was monitored. After the heat shock, plants were cooled down at room temperature for 30 minutes and grown in the growth chamber for additional 3 to 4 weeks until the lamina joint distance between the flag leaf and a leaf below reached 3 to 5 cm. At this stage, the length of internode II ranged from 1 to 8 cm.

### RNA extraction, cDNA synthesis, and RT-PCR

RNA extraction, cDNA synthesis, reverse transcriptase polymerase chain reaction (RT-PCR), and quantitative RT-PCR (qRT-PCR) were performed as described previously (Tsuda et al., 2022). Total RNA was extracted from germinating seedlings at 3 DAG. At least three plants were pooled in each biological replicate, and three biological replicates were performed for each data point. Relative expression levels were calculated using the 2^–ΔΔCt^ method, and the rice ubiquitin (*ubq*) gene was used as an internal standard. Primers used in PCR reactions are listed in **Supplemental Tabel 4**.

### Recording and representation of GUS sectors in the flag leaf phytomer

GUS staining was conducted as described previously (Jefferson et al., 1987; Tsuda et al., 2022). The stained sectors were observed under the dissecting microscope (SZX16, Olympus). We dealt with only epidermal and subepidermal sectors that can be unambiguously traced. To record the approximate positions of sectors, each domain (foot II, internode II, node I, the pulvinus, and the flag leaf sheath) was divided into three bins: bottom, middle, and top thirds. The relative position of each sector was recorded in spreadsheets, in which rows and columns correspond to samples and bins, respectively. After rows in this spreadsheet were sorted by the relative position of sectors, data were plotted using the “heatmap” function in base R.

### Micro-computed tomography (micoro-CT)

Micro-CT scanning was performed as described previously (Tsuda et al., 2017; Maeno and Tsuda, 2018). Rice stem samples (5 mm) were fixed in FAA (formalin:acetic acid:50% ethanol = 5:5:90) overnight. The fixative was replaced with 70% ethanol and stored at 4°C until observation. Before scanning, samples were soaked in the contrast agent (0.3% phosphotungstic acid in 70% ethanol) for 18 days and scanned using x-ray micro-CT at a tube voltage peak of 80 kVp and a tube current of 90 µA. Samples were rotated 360° in steps of 0.24°, generating 1500 projection images of 992 x 992 pixels. The micro-CT data were reconstructed at an isotropic resolution of 5.5 x 5.5 x 5.5 µm^3^. Three-dimensional tomographic images were obtained using the OsiriX software.

### Histological observation

Shoot apices were dissected and fixed in FAA (formalin: acetic acid: ethanol: water = 10:5:50:35) and were embedded in Paraplast Plus (McCormick Scientific) as described previously (Tsuda et al., 2017). Eight µm sections were stained using toluidine blue O and observed under light microscopy (Olympus BX50). To count epidermal cells in the developing stem, sections were stained with calcofluor white and imaged under the Fluoview FV300 CLSM system (Olympus).

## Supplemental data

Supplemental Figure 1: Histological observations of stem development during initiation of the flag leaf phytomer.

Supplemental Table 1: GUS sector induction rate in the vegetative shoot apex.

Supplemental Table 2: GUS sector induction rate in the internode II.

Supplemental Table 3: The details of sectors extended for multiple tissue types (TT). Supplemental Table 4: Primers used in this study.

## Author contributions

K.T. designed this work, conducted experiments, and analyzed data. A.M. performed the micro-CT observation. K.I.N. supervised the project. K.T. wrote the manuscript with the help of K.I.N.

## Acknowledgments

We thank Ari Pekka Mähönen for providing original binary vectors for the clonal analysis in *Arabidopsis* and Keisuke Nagai for critical reading and comments on the manuscript. We also thank Kae Kato and Ayako Otake for their technical assistance. This work is supported by JSPS KAKENHI 19K05980, 20H04891, 22H02319 and 23H04754 to K.T, and 21H04729 to K.I.N.

